# Relief from nitrogen starvation entails quick unexpected down-regulation of glycolytic/lipid metabolism genes in enological *Saccharomyces cerevisiae*

**DOI:** 10.1101/428979

**Authors:** Tesnière Catherine, Bessière Chloé, Pradal Martine, Sanchez Isabelle, Blondin Bruno, Bigey Frédéric

## Abstract

Nitrogen composition of the grape must has an impact on yeast growth and fermentation kinetics as well as on the organoleptic properties of the final product. In some technological processes, such as white wine/rosé winemaking, the yeast-assimilable nitrogen content is sometimes insufficient to cover yeast requirements, which can lead to slow or sluggish fermentations. Growth is nevertheless quickly restored upon relief from nutrient starvation, e.g. through the addition of ammonium nitrogen, allowing fermentation completion. The aim of this study was to determine how nitrogen repletion affected the transcriptional response of a *Saccharomyces cerevisiae* wine yeast strain, in particular within the first hour after nitrogen addition. We found almost 4800 genes induced or repressed, sometimes within minutes after nutrient changes. Some of these responses to nitrogen depended on the TOR pathway, which controls positively ribosomal protein genes, amino acid and purine biosynthesis or amino acid permease genes and negatively stress-response genes, and genes related to the retrograde response (RTG) specific to the tricarboxylic acid (TCA) cycle and nitrogen catabolite repression (NCR). Some unexpected transcriptional responses concerned all the glycolytic genes, carbohydrate metabolism and TCA cycle-related genes that were down-regulated, as well as genes from the lipid metabolism.

## Introduction

The yeast cell *Saccharomyces cerevisiae* is able to control its growth in response to changes in nutrient availability. Nitrogen limitation is one of the most frequent limitations observed during wine fermentation [1]. The actual nitrogen content in must is dependent on many factors including rootstock, grape variety, climate, vine growing conditions, and grape processing. In enological conditions, musts are considered as nitrogen-limited when the yeast assimilable nitrogen (YAN) content is below 150 mg/L [1]. YAN is a major factor influencing fermentation kinetics, the maximal fermentative rate being related to the nitrogen level in the must [1]. In most cases of sluggish fermentations, nitrogen depletion quickly results in cells entering stationary phase. This phenomenon is not related to a decrease in viability, but could rather be related to a catabolic inactivation of the hexose transporters [2] or to lower protein synthesis and cell protein content [3]. Other physiological changes such as autophagy, nitrogen recycling systems and the reorientation of the carbon flux to promote glycogen and trehalose storage have also been observed at the onset of nitrogen starvation [4]. In addition, the transcriptional remodeling associated with the onset of starvation during wine alcoholic fermentations has been described [3], including the development of a general stress response. These transcriptional changes are mostly controlled by the TOR pathway, sensing cell nitrogen status and adapting nitrogen metabolism to nutrient availability [5, 6]. Nitrogen limitation stably arrests the cell cycle in G_1_/G_0_, whereas medium replenishment with the limiting nutrient quickly restores growth. Relief from nitrogen starvation is a way to increase the fermentation rate, while reducing its duration [7]. In fact assimilable nitrogen addition to nitrogen-deficient must results in a reactivating protein synthesis and increasing sugar transport speed [7, 8]. Although this nitrogen addition is currently practiced using diammonium phosphate to reduce the risk of stuck fermentation in white and rosé wines, the molecular mechanisms triggered by nitrogen replenishment are still poorly understood.

The present work complements previous investigations on laboratory [9] or enological [10] yeast strains with the novelty of transcriptome analysis every 15 min during the first hour following relief of nitrogen starvation in a medium mimicking grape must composition with limiting nitrogen concentration. We report here rapid transcriptional changes that occur in a wine yeast strain in response to relief from nitrogen starvation. Our goal was to detect new phenomena appearing quickly after nitrogen addition.

## Materials and methods

The experimental design was mapped out on S1 Fig.

### Strain and culture conditions

All fermentation experiments were carried out in triplicates (S1 Fig) using the yeast strain *Saccharomyces cerevisiae* Lalvin EC1118, a commercial wine yeast from Lallemand Inc (Canada). The culture medium was a synthetic medium [1] that mimics a standard natural must. In our conditions the total concentration of yeast assimilable nitrogen (YAN) was 100 mg/L and we added 24.1 mg/L FeCl_3_ · 6H_2_O (see SI Experimental Procedures). Fermentations were conducted in 1 L of medium under constant stirring at 24 °C. Flasks (1.2 L) were equipped with locks to maintain anaerobiosis. Production of CO_2_ was monitored by weighing the flasks every 20 min, to determine weight loss. The rate of CO_2_ production was estimated using a polynomial smoothing as previously described [11]. The number of cells was determined with a particle counter (Coulter counter, Beckman Coulter). Preliminary experiments have shown that, under this condition, cells were starved for nitrogen (i.e. reached stationary phase) after 42 h when 14 g of CO_2_ has been released [7, 12]. Some cells were collected at this stage as controls (*t* = 0), then diammonium phosphate (DAP, (NH_4_)_2_HPO_4_) was added to the culture medium (300 mg/L final concentration), after removing an equivalent volume of medium to keep the total volume unchanged. This supplement provides 63 mg/L of atomic nitrogen, entirely assimilable, corresponding to the maximum nitrogen addition permitted in wine-making.

Sampling was then performed 15, 30, 45 and 60 min after DAP addition and cells were quickly recovered by filtration and frozen at −80 °C as previously described [9].

### Labeling and microarray processing

Total RNA extraction was performed with Trizol reagent, and purified with RNeasy kit (Qiagen). Spike-in RNAs were added to 100 ng total RNA using the One-color RNA Spike-In kit (Agilent Technologies) and Cy3-labeled cRNAs were synthesized using the Low Input Quick Amp Labeling kit (one-color, Agilent Technologies). Labeled probes were purified with RNeasy kit (Qiagen). Quality and quantity of RNA were controlled at each step by spectrometry (NanoDrop 1000, Thermo Scientific). Labeled cRNA were hybridized to custom 8×15K microarray (Agilent Technologies) containing the Yeast V2 probe-set (Agilent ID: 016322) together with 39 probes corresponding to *Saccharomyces cerevisiae* EC1118 specific genes [13]. This design was registered in the Gene Expression Omnibus (GEO) repository under platform accession number GPL17690. 600 ng of labeled cRNA were hybridized for 17 h at 65 °C in a rotative hybridization oven (Corning) using the Gene Expression Hybridization kit (Agilent Technologies). Array digitalization was performed on a GenePix^®^ 4000B laser Scanner (Axon Instruments) using GenePix^®^ Pro7 Microarray Acquisition and Analysis Software (Axon Instruments). Data normalization and statistical analysis were performed using R software [14] and the limma package [15]. Normalization was performed by the quantile method considering all arrays. The resulting absolute expression levels were expressed as logarithm (base 2) for each time and replicate. The data were deposited in GEO under accession number GSE116766 (also available in S1 Table).

### Statistical analysis

Normalized data were first converted to fold changes relative to expression at *t* = 0, then we analyzed changes over time using a regression based approach to find genes with temporal expression changes (S2 Fig). We defined a binomial regression model for each gene expression over 5 time points: *Y* = *b*_0_ + *b*_1_*t* + *b*_2_*t*^2^ + *ϵ*, where *Y* is the normalized expression value, *t* is the time (min), *b*_0_ is expression at *t* = 0, *b*_1_ is the slope (induction or repression of the gene, linear effect), *b*_2_ is a quadratic effect and *ϵ* is the residual error term. A variable selection procedure was applied using step regression (backward method) to find significant coefficients for each gene. We adjusted this model by the least-squared technique for each gene and only genes with significant changes over time were selected with an adjusted p-value threshold of 0.01 corrected by the Benjamini-Hochberg method. Distribution of *b*_1_ and *b*_2_ coefficients is presented on S2 Fig. The sign of *b*_1_ distinguish between up (positive, clusters 1,3,5,7) and down-regulated (negative, clusters 2,4,6,8) gene expression. Furthermore, the sign of *b*_2_ allow us to distinguish between accelerated (positive, clusters 2,5,7) and decelerated (negative, clusters 1,6,8) expression rate. Genes belonging to clusters 3 and 4 (*b*_2_ = 0) have linear expression profiles.

Functional analysis was performed looking for Gene Ontology (GO) term enrichment (biological process) using GO Term Finder [16] with the multiple test correction of Benjamini Hochberg.

## Results and Discussion

### Changes in fermentation kinetics after nitrogen repletion

We investigated the very early events occurring after relief of nitrogen starvation in a wine strain under enological conditions, by samplings every 15 min during the first hour of replenishment. Fig 1 presents a typical fermentation kinetics in a nitrogen-limited synthetic must [1]. First a rapid increase of the CO_2_ production rate was observed, reaching a maximum (0.9 g/L/h) at 25 h after inoculation. Thereafter, the rate decreased sharply indicating an arrest of the population growth, the so called stationary phase, where nitrogen was limiting beginning at 42h (14 g of CO_2_ released). Then the production rate decreased slowly up to 280 h (corresponding to 93 g of CO_2_ released), indicating that all glucose had been converted to CO_2_ and ethanol. If diammonium phosphate (DAP) was added at the beginning of the stationary phase (42 h), a very quick restart of the rate of CO_2_ production which peaked (1.2 g/L/h) higher than the maximum reached at the beginning of the fermentation (0.7 g/L/h). Fermentation ended in 190 h, reducing the fermentation duration by almost 30%. As previously described, DAP addition to nitrogen-starved wine yeast cells resulted in a very quick restart of the rate of CO_2_ production [7, 17]. During the course of the sampling experiment (every 15 min for 60 min after DAP addition), nitrogen is not expected to be limiting as it was found that nitrogen was completely consumed only after 4 hours under the same conditions [17].

**Fig 1.**
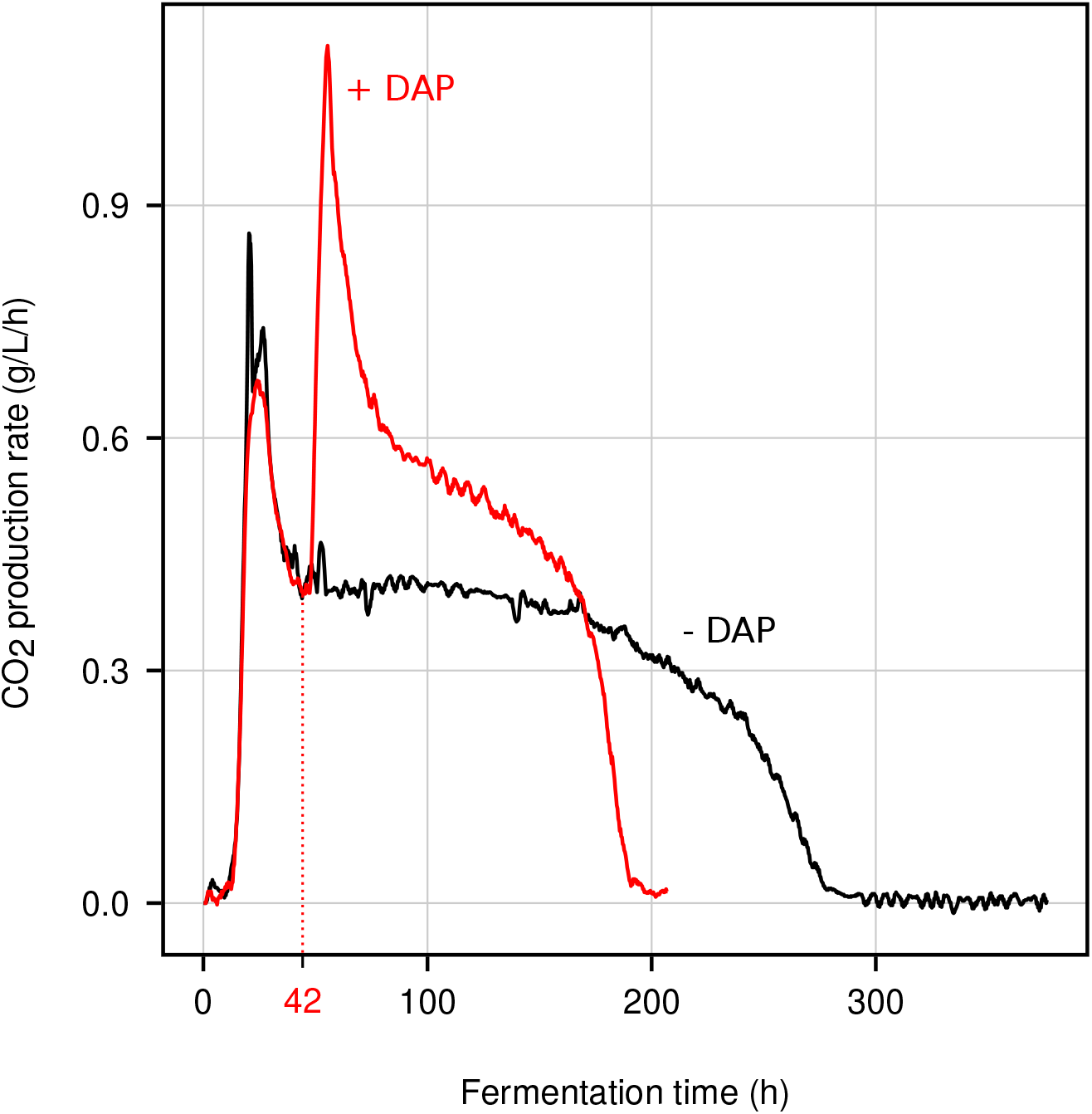
**Fermentation profiles CO_2_** production rate during fermentation in a nitrogen-depleted synthetic must (black). In another experiment (red), DAP was added at the beginning of the stationary phase (42 h; 14 g CO_2_ released)

### Numerous changes in gene expression

#### Significantly regulated genes

We studied the expression of yeast genes within 1 hour following DAP addition at 0, 15, 30, 45 and 60 min. Step regression analysis revealed numerous changes during this first hour with almost 4800 nitrogen-regulated genes identified (S2 Table). This is much higher than the 350 genes regulated after 2 hours upon the addition of DAP to active dried yeast inoculated in a Riesling must [18], or than the 1000 [19] or 3000 [9] transcripts altered by the addition of nitrogen to laboratory yeast cells. These differences are probably due to improvements in the DNA microarray technology, to a reduced time-scale or to the experimental conditions (synthetic versus natural must, industrial versus laboratory yeast strains).

Thereafter, genes were classified using manual clustering (S2 Fig) in 8 expression profiles (S2 Table). Respectively 2292 (clusters 1, 3, 5, 7; Fig 2) and 2507 (clusters 2, 4, 6, 8; Fig 3) genes were significantly up- or down-regulated, reflecting a massive change in expression patterns upon nitrogen repletion. For each cluster, individual gene expression is available in S3 Table.

**Fig 2.**
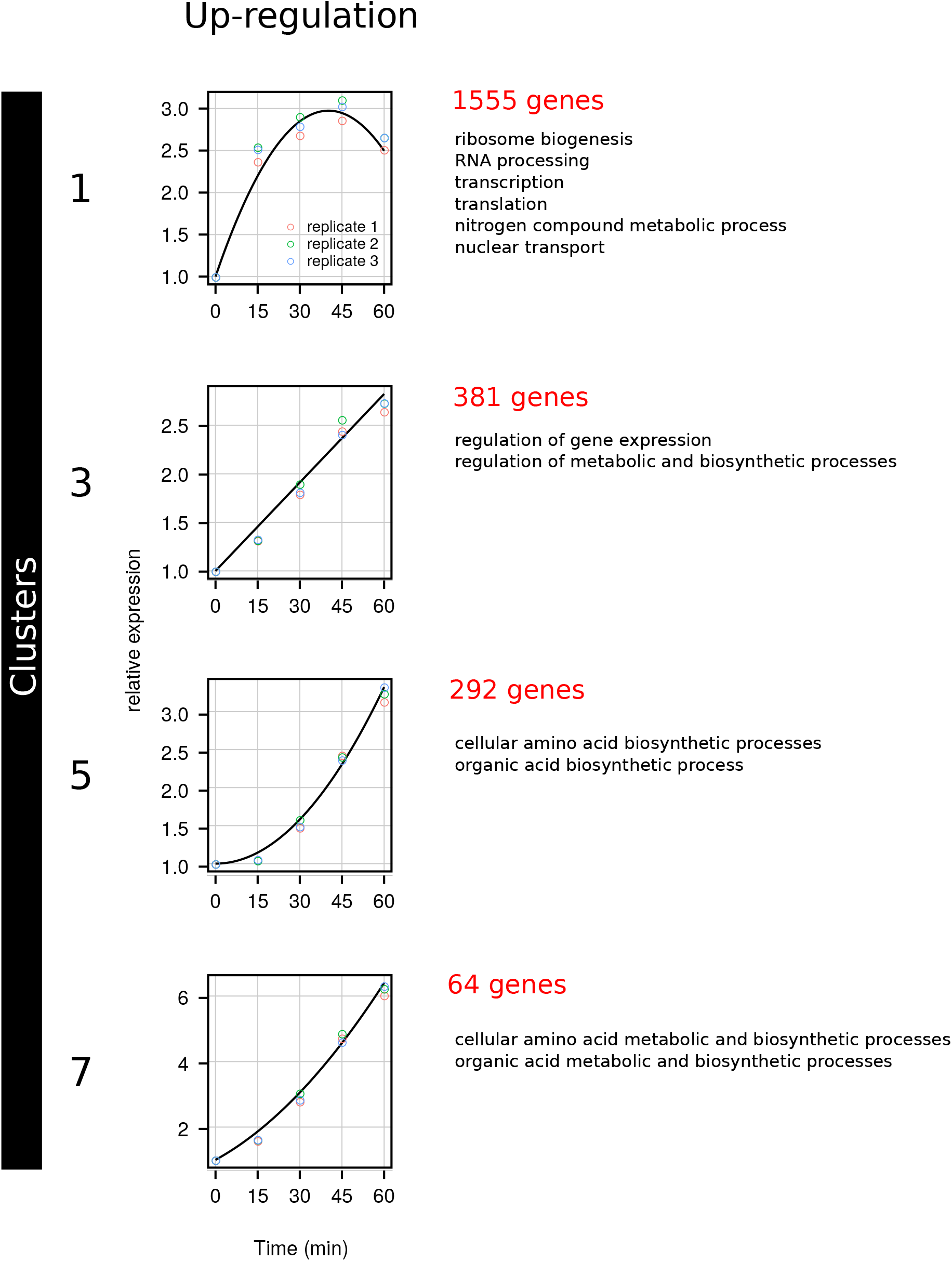
Clusters of up-regulated genes. Clustering of expression pattern and GO-term enrichment were performed as described in the Materials and Methods

**Fig 3.**
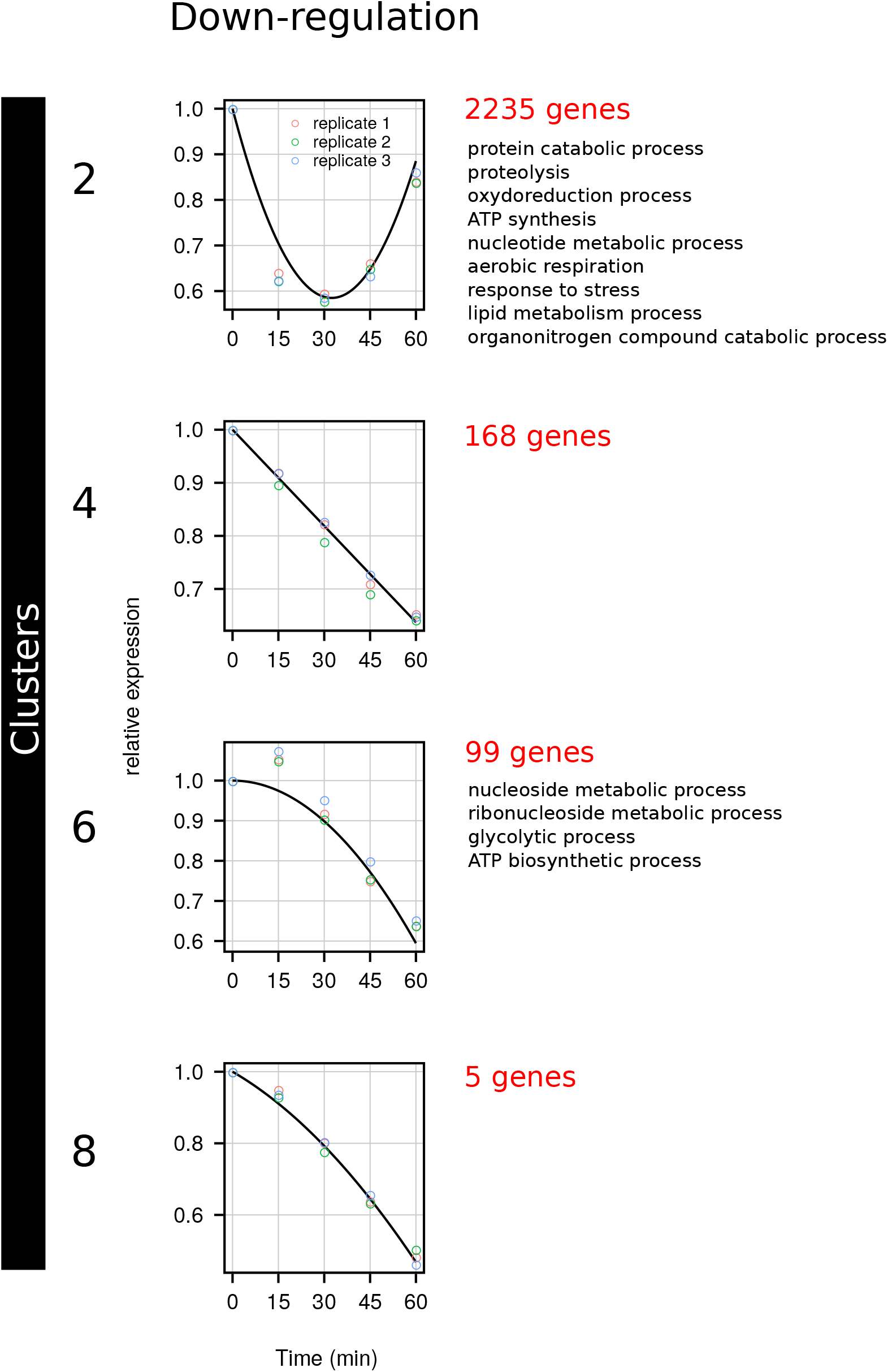
Clusters of down-regulated genes. Clustering of expression pattern and GO-term enrichment were performed as described in the Materials and Methods

#### Up-regulated genes

Among the clusters corresponding to up-regulated genes (Fig 2), cluster 1 contained 1555 genes exhibiting an initial linear increase (*b*_1_ > 0, S2 Fig), sharp but transient, then a decrease of expression due to the negative quadratic term of the equation (*b*_2_ < 0). Functional analysis using GO-term enrichment (S4 Table) showed that this cluster contained many genes involved in or related to ribosome biogenesis, RNA processing, transcription, translation, nitrogen compound metabolic process, nuclear transport. Cluster 3 contained 381 genes linearly induced within the first hour following repletion (*b*_1_ > 0 and *b*_2_ =0), encoding proteins involved in the regulation of gene expression and of metabolic and biosynthetic processes. Cluster 5, which contained 292 genes that exhibited an expression accelerating with time (*b*_1_ = 0 and *b*_2_ > 0), was enriched in genes involved in amino-acid (*TRP2, MET8, HIS3, LEU4, TRP3, LYS2, HIS5, ARG4, HIS4, ARG7, ARG1*) and organic acids biosynthetic processes. Cluster 7 contained 64 genes that exhibited the highest increase in expression among all up-regulated genes, following a linear profile (*b*_1_ > 0) with a slight acceleration (*b*_2_ > 0). Functional analysis showed that this cluster was similar to cluster 5.

This global response is similar to what was described for the commercial wine yeast strain VIN13, 2 hours after DAP addition [18], where an up-regulation was observed for genes involved in amino acid metabolism, de novo purine biosynthesis, and protein synthesis. Such changes likely corresponded to an activation of the Target of Rapamycin (TOR) signaling pathway which positively controls ribosomal protein genes [20], amino acid and purine biosynthesis or amino acid permease genes [21]. Surprisingly, within 60 min we didn’t find any change in the expression of genes related to sulfate assimilation, although this had been observed (after two hours) by [18]. This is probably due to the fact that the authors used true grape must instead of synthetic grape must, which resulted in a difference in concentration between sulfur-containing compounds, methionine and cysteine.

Three components of the MCM (mini-chromosome maintenance) hexameric complex helicase, binding to chromatin as a part of the pre-replicative complex (*MCM2, MCM3*, and *MCM6*), and also *MAD1* and *YCG1*, were transiently but sharply induced after relief from nitrogen starvation. The MCM complex is required for the initiation of eukaryotic replication, while Mad1p is a coiled-coil protein involved in the spindle-assembly checkpoint. Its phosphorylation leads to an inhibition of the activity of the anaphase promoting complex. Ycg1p is required for establishment and maintenance of chromosome condensation, chromosome segregation and chromatin binding of the condensin complex and is also required for clustering tRNA genes at the nucleolus. In addition, other cell-cycle related genes were induced, such as *CLN3, SWI6, RAD59, CDC20, RFA3, MSH2* and *YHM2*. Thus, all these transient inductions are coherent with a restart of the cell cycle in response to nitrogen replenishment.

#### Down-regulated genes

Among the clusters corresponding to down-regulated genes (Fig 3), 2235 genes in cluster 2 exhibited an initial linear decrease (*b*_1_ < 0), sharp but transient, then an increase of expression due to the positive quadratic term of the equation (*b*_2_ > 0). Functional analysis (S4 Table) showed that cluster 2 contained many genes involved in protein catabolic process, proteolysis, organonitrogen compound catabolic process, lipid metabolic process, response to stress, oxido-reduction process, ATP synthesis, nucleotide metabolic process and aerobic respiration. Cluster 4 contained 168 genes that were linearly repressed during the first hour following repletion (*b*_1_ < 0 and *b*_2_ = 0). No significant enrichment in GO-terms was observed for this cluster. Cluster 6 contained 99 genes that exhibited a decelerating expression with time (*b*_1_ = 0 and *b*_2_ < 0) and was enriched in genes involved in nucleoside and ribonucleoside metabolic process, glycolytic process and ATP biosynthetic process. Finally, cluster 8 contained only 5 genes that exhibited an amplitude of down-regulation similar to the previous clusters. This is a linear profile (*b*_1_ < 0) with a slightly deceleration (*b*_2_ < 0). Functional analysis showed no significant enrichment.

In our conditions (i.e. within one hour after repletion), we found other functions for down-regulated genes than those described previously [18]. In fact, genes related to cellular transport were repressed in response to DAP addition (*NCE102, POR1, PMA2, ATP19, ATP2, UGA4, PUT4, GSP2, YPT53*). Other most interesting genes were those related to stress response, those sensitive to the nitrogen catabolite repression (NCR), and those related to the glycolysis. Another group of genes are related to lipid biosynthesis. Among this last group, we found *ETR1, IFA38, ERG28, ERG4, ERG25, ERG11, NCP1, ERG20, ELO1, FAS1, ERG3, ERG6, ERG5, LIP1, ERG24, ACC1, POT1, TIP1, OPI3, YML131W, AAD10, GCY1, GRE3, TGL4* and, more specifically those related to ergosterol biosynthesis (*ERG28, ERG4, ERG25, ERG11, NCP1, ERG20, MCR1, ERG3, ERG6, ERG5, ERG24, ERG10*). This discovery could be explained by the fact that the biosynthesis of lipids requires a lot of energy, unavailable at the resumption of fermentation when the biosynthesis of proteins increases significantly.

Moreover, DAP addition decreased the expression of a large group of genes of the Ras/Protein Kinase A (PKA) signaling pathway (*IRA1, IRA2, GPR1, GPA2, CYR1, TPK1, TPK2, BCY1, SCH9, YAK1*) and genes related to the stress response, such as genes coding the heat-shock proteins, but also genes related to the seripauperin multigene family (PAU), which mostly belong to cluster 2. This pattern indicated that the down-regulation of these genes was a rapid phenomena, largely decreasing within the first 15 min after nitrogen repletion. Other genes related to stress gene regulation such as *HSF1, MSN2*, and *MSN4* [22] were also down-regulated in our study as well as genes involved in trehalose and glycogen metabolisms (*TPS1, TPS2, TPS3, ATH1, NTH1, NTH2, TSL1, GPH1, GPD1, GSY1, GSY2*).

Such changes are also likely related to an activation of the TOR signaling pathway that also negatively controls stress-response genes, the retrograde response (RTG) specific to the tricarboxylic acid (TCA) cycle genes and genes sensitive to the nitrogen catabolite repression (NCR) [21].

Interestingly, the down-regulation of genes related to glycolysis, which has been previously reported in similar experimental conditions but on a laboratory strain [9], was confirmed here in an enological strain (Fig 4). This indicates the conservation of this mechanism independently of the yeast strain used. As previously suggested, these unexpected results were probably revealed by analyzing the very early events following nitrogen replenishment. This repression of glycolytic genes in wine yeast had already been observed, but in rather different experimental conditions, such as 1 h after inoculation of a synthetic must [10]. It has been hypothesized that this destabilization of transcripts know to be stable might be a consequence of the recovery of protein synthesis upon addition of nitrogen on starved yeasts [9].

**Fig 4.**
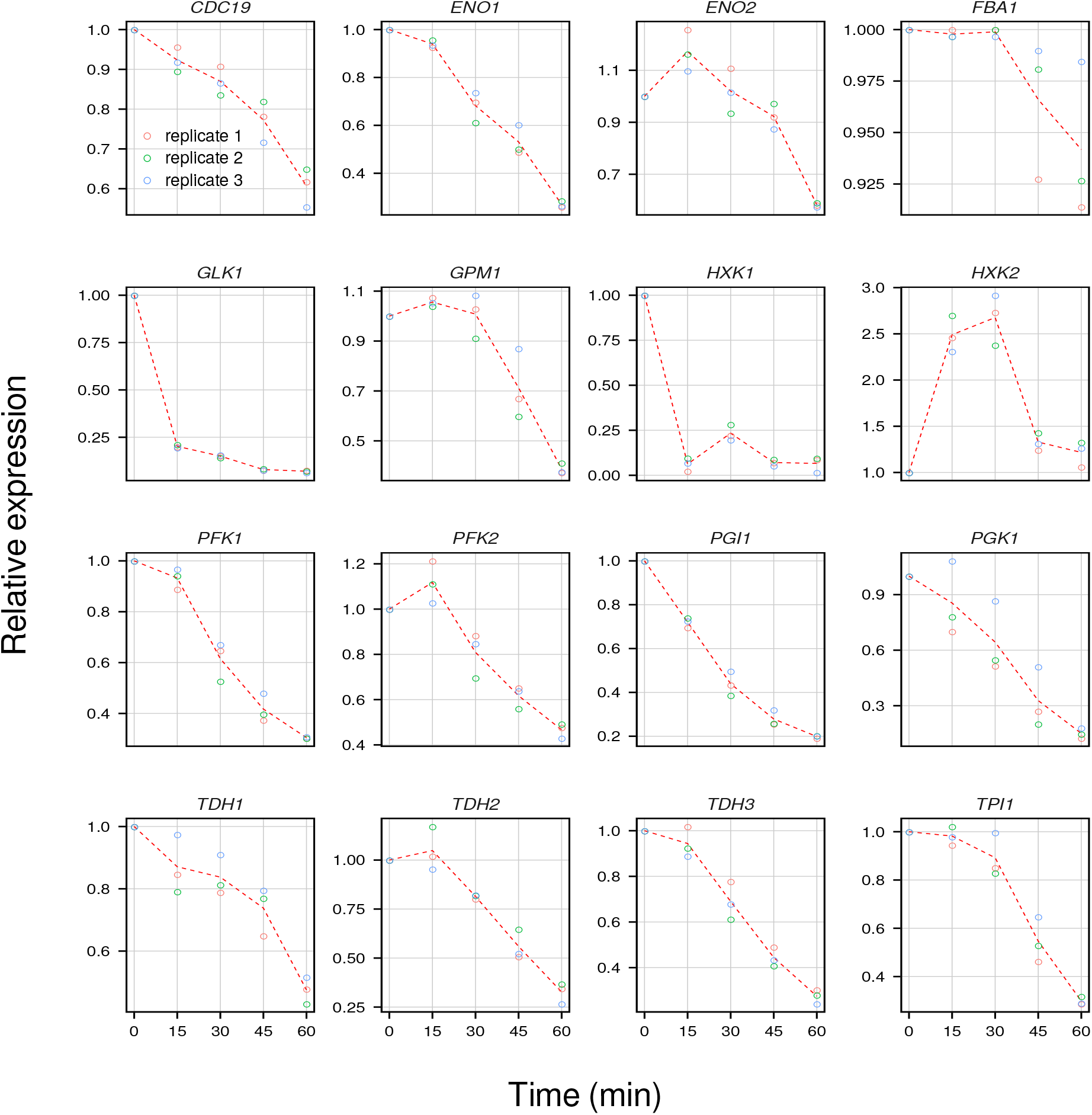
Expression profiles of glycolytic genes. Expression profile of 16 glycolytic genes

Other important changes were also revealed, in the present study, concerning for instance the down-regulation of genes related to the MAPK signaling pathways, oxidoreductase activity, or sensitive to NCR. Concerning genes related to stress and NCR, their down-regulation corresponded to a common response to glucose, nitrogen and phosphorous repletion, whereas the down-regulation of nitrogenous compound catabolism and amino acid derivative transport were nitrogen-specific [19]. For these authors, both PKA and TOR signaling pathways might be involved in the responses to all three nutriments viz. glucose, nitrogen and phosphate. Surprisingly, these authors found that genes associated with glycolysis and gluconeogenesis were specifically repressed by phosphorous, whereas in the present study they were both nitrogen- and phosphate-regulated (as we used ammonium phosphate).

It was in fact surprising to observe the repression of all the glycolysis-related genes whereas genes related to ribosomal protein synthesis were up-regulated. This could indicate that the restart of the fermentative activity shortly after the addition of DAP was unrelated to the glycolytic pathway but rather to the cell cycle and protein synthesis activation. In fact, Rim15p, which gene expression is down-regulated in our study, has been found to integrate signals derived from PKA, TORC1 and Sch9p, which transmit the information concerning the availability of nutrients [23]. Rim15p regulates proper entry into G_0_ via the transcription factors Msn2/4p and Gis1p whose related genes were also down-regulated. The down-regulation of *RIM15* is thus coherent with the up-regulation of cell-cycle related genes and correspond to the model previously suggested [9].

## Conclusion

The addition of nitrogen to starved wine yeast cells thus contributed to the development of a favorable environment for wine yeast growth and also to limit the general stress response. During a very short time after the addition of nitrogen to the medium, we found thousands of genes induced or repressed, sometimes within minutes after nutrient changes. Some of these responses to nitrogen depended on the TOR pathway, which controls positively ribosomal protein genes, amino acid and purine biosynthesis or amino acid permease genes and negatively stress-response genes, and genes related to the retrograde response (RTG) specific to the tricarboxylic acid (TCA) cycle and nitrogen catabolite repression (NCR). Most of these responses are the opposite of the changes observed in yeasts deprived of nitrogen, when the cells reach the stage of the stationary phase [4]. But we also detected unexpected transcriptional responses. These included all glycolytic genes, carbohydrate metabolism and TCA cycle genes that were downregulated, as well as genes derived from lipid metabolism.

## Supporting information

SI Supplemental procedures

## Supporting information

**SI Experimental Procedures. Supplementary experimental procedures**. Additional materials and procedures. (PDF)

**S1 Fig. Experimental design**. Schematic representation of the experimental design. (PDF)

**S2 Fig. Statistical analysis methods**. Schematic representation of methods used to analyze the expression data: selection of a model for the time-course experiment, step-regression and manual clustering of expression profiles. (PDF)

**S1 Table. Raw gene expression after normalization**. For each replicate at each time point, this table gives the absolute expression level as expressed in logarithm to the base 2. (TSV)

**S2 Table. Step regression and clustering result**. This spreadsheet contains the regression coefficients and the statistical supports obtained after step regression for each regulated genes. (XLSX)

**S3 Table. Clusters’ composition**. This spreadsheet presents the gene-composition of each cluster. For each gene at each time point, the expression levels are expressed relative to that at *t*_0_. (XLSX)

**S4 Table. Functional analysis**. This spreadsheet contains the GO-term enrichment for each cluster. (XLSX)

## Acknowledgments

We wish to warmly acknowledge Dr Philippe Chatelet for his useful suggestions on the text. We would like to thank reviewers for their insightful comments.

## Author contributions

C.T., B.B. and F.B. jointly conceived the study, interpreted the data and wrote the paper. C.B. and M.P. performed experiments. I.S. and F.B. conducted statistical analyses of microarray data.

## Data availability

Dataset, R scripts, figures and tables are available in open-access on Zenodo (doi:10.5281/zenodo.1295508)

