## Supplementary material for "Relief from nitrogen starvation entails quick unexpected down-regulation of glycolytic/lipid metabolism genes in enological *Saccharomyces cerevisiae*": SI Supplemental procedures

### SI Experimental Procedures

Frédéric Bigey

March 21, 2019

### 1 Synthetic must composition

The fermentation medium mimics a standard natural must and has the same composition as the synthetic medium previously described [1], except for the total concentration of assimilable nitrogen that was 100 mg/L and the addition of 24.1 mg/L $FeCl_3 \cdot 6\,H_2O$ (Table 1). Chemicals were mixed in 300 mL of warm deionized (DI) water, volume was adjusted to 950 mL with DI water and pH adjusted to 3.3 with NaOH (12N). Finally, the volume was adjusted to 1 L with DI water.

| Chemical | Quantity for 1 L |
|---|---|
| glucose | 200 g |
| DL malic acid | 6 g |
| citric acid | 6 g |
| $KH_2PO_4$ | 0.75 g |
| $K_2SO_4$ | 0.5 g |
| $MgSO_4 \cdot 7\,H_2O$ | 0.25 g |
| $CaCl_2 \cdot 2\,H_2O$ | 0.155 g |
| NaCl | 0.20 g |
| $NH_4Cl$ | 0.46 g |
| amino-acids solution (Table 2) | 3.08 mL |
| oligo-elements solution (Table 3) | 1.00 mL |
| vitamins solution (Table 4) | 10.00 mL |
| lipid & sterol solution (Table 5) | 1.00 mL |
| $FeCl_3 \cdot 6\,H_2O$ (17 g/L) | 1.42 mL |

Table 1: Must composition

| Chemical | Quantity for 1 L |
|---|---|
| $NaHCO_3$ | 20.0 g |
| aspartic acid | 3.4 g |
| glutamic acid | 9.4 g |
| alanine | 11.1 g |
| arginine | 28.6 g |
| cysteine | 1.1 g |
| glutamine | 38.6 g |
| glycine | 1.4 g |
| histidine | 2.5 g |
| isoleucine | 2.5 g |
| leucine | 3.7 g |
| lysine | 1.3 g |
| methionine | 2.4 g |
| phenylalanine | 2.9 g |
| proline | 46.8 g |
| serine | 6.0 g |
| threonine | 5.8 g |
| thryptophan | 13.7 g |
| tyrosine | 1.4 g |
| valine | 3.4 g |

Table 2: Amino-acids solution: In 400 mL of hot DI water, dissolve 20 g of $NaHCO_3$ then add amino-acids. Let solution cool down to room temperature then adjust volume to 1 L with DI water. Store solution at $-20$ °C.

| Chemical | Quantity for 1 L |
|---|---|
| $MnSO_4 \cdot H_2O$ | 4.0 g |
| $ZnSO_4 \cdot 7\,H_2O$ | 4.0 g |
| $CuSO_4 \cdot 5\,H_2O$ | 1.0 g |
| KI | 1.0 g |
| $CoCl_2 \cdot 5\,H_2O$ | 0.4 g |
| $H_3BO_4$ | 1.0 g |
| $(NH_4)_6Mo_7O_{24}$ | 1.0 g |

Table 3: Oligo-elements solution: In 300 mL of DI water, dissolve the chemicals then adjust volume to 1 L with DI water. Filter-sterilize and store solution at +4 °C.

| Chemical | Quantity for 1 L |
|---|---|
| myo-inositol | 2.0 g |
| calcium pantothenate | 0.15 g |
| thiamine hydrochloride | 0.025 g |
| nicotinic acid | 0.2 g |
| pyridoxine | 0.025 g |
| biotine (100 mg/L) | 3 mL |

Table 4: Vitamins solution: In 300 mL of DI water, dissolve the vitamins then adjust volume to 1 L with DI water. Store solution at −20 °C.

| Chemical | Quantity for 100 mL |
|---|---|
| ergosterol | 1.5 g |
| oleic acid | 0.5 mL |
| Tween 80 | 50 ml |
| ethanol | to 100 mL |

Table 5: Lipids and sterols solution: Dissolve ergosterol and oleic acid in 100 mL of a mixture of Tween 80/ethanol (50/50, v/v). Warm the mixture at 70 °C for 5 min with mixing. Store the mixture at +4 °C.
