## Supplementary figures and images for "Relief from nitrogen starvation entails quick unexpected down-regulation of glycolytic/lipid metabolism genes in enological *Saccharomyces cerevisiae*"

### S1 Fig

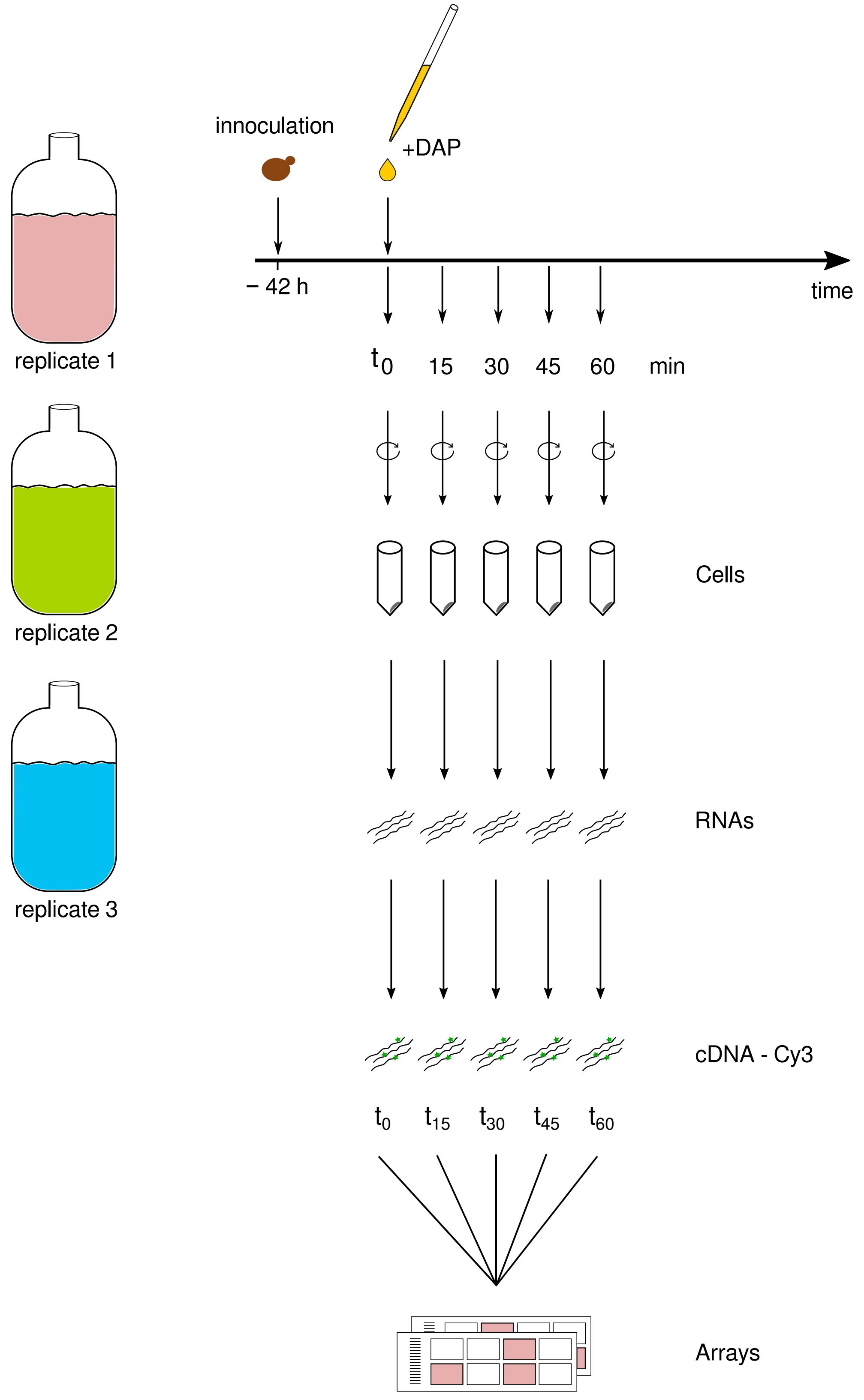

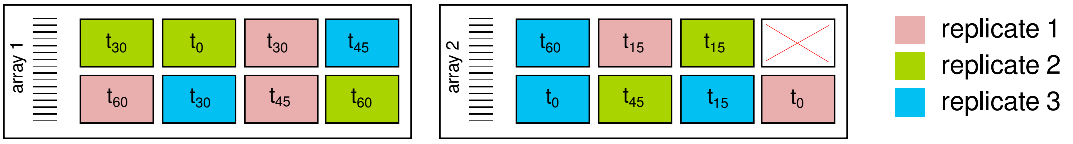
