## Supplementary material for "Relief from nitrogen starvation entails quick unexpected down-regulation of glycolytic/lipid metabolism genes in enological *Saccharomyces cerevisiae*": S2 Fig

### S2 Figure

$Y = b_0 + b_1t + b_2t^2 + \varepsilon$

model of gene expression over time

**Step 1**

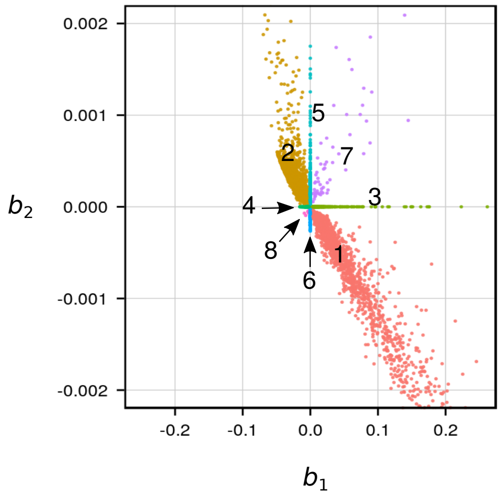


estimation of model coefficients by step regression

cluster identification

**Step 2**

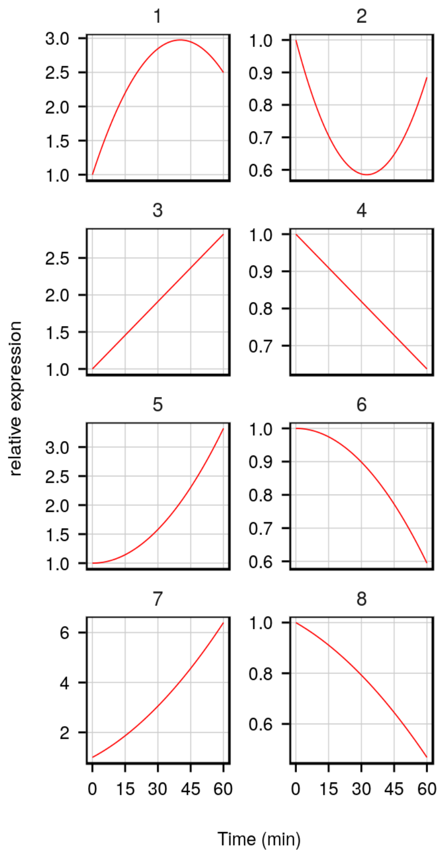


average expression per cluster

**Step 3**
